# Role of the novel endoribonuclease SLFN14 and its disease causing mutations in ribosomal degradation

**DOI:** 10.1101/267633

**Authors:** Sarah J. Fletcher, Vera P. Pisareva, Abdullah Khan, Andrew Tcherepanov, Neil V. Morgan, Andrey V. Pisarev

## Abstract

Platelets are anucleate and mostly ribosome-free cells within the bloodstream, derived from megakaryocytes within bone marrow and crucial for cessation of bleeding at sites of injury. Inherited thrombocytopenias are a group of disorders characterized by alow platelet count and are frequently associated with excessive bleeding. *SLFN14* is one of the most recently discovered genes linked to inherited thrombocytopenia where several heterozygous missense mutations in *SLFN14* were identified to cause defective megakaryocyte maturation and platelet dysfunction. Yet, SLFN14 was recently described as a ribosome-associated protein resulting in rRNA and ribosome-bound mRNA degradation in rabbit reticulocytes. To unveil the cellular function of SLFN14 and the link between SLFN14 and thrombocytopenia, we examined SLFN14 (WT/mutants) in in vitro models. Here, we show that all SLFN14 variants co-localize with ribosomes and mediate rRNA endonucleolytic degradation and ribosome clearance. Compare dto SLFN14 WT, expression of mutants is dramatically reduced as a result of post-translational degradation due to partial misfolding of the protein. Moreover, all SLFN14 variants tend to form oligomers. These findings could explain the dominant negative effect of heterozygous mutation on SLFN14 expression in patients’ platelets. Overall we suggest that SLFN14 could be involved in ribosome degradation during platelet formation and maturation.

## INTRODUCTION

Inherited thrombocytopenias (ITs) are a group of disorders determined by a relative decrease of platelet count resulting from a genetic heterogeneity [Nurden and Nurden 2007]. ITs are usually asymptomatic, but some individuals may experience excessive bleeding ranging from mild to severe. Over 30 genes are shown to be involved in ITs [Favier and Raslova 2015; Levin et al. 2015; Johnson et al. 2016a; Johnson et al. 2016b; Pecci 2016]. One of these, *SLFN14*, was discovered only very recently where Fletcher et al. identified three heterozygous missense mutations in affected family members from 3 unrelated families, predicted to encode substitutions K218E, K219N, and V220D within an AAA domain of SLFN14 [Fletcher et al. 2015]. Patients revealed moderate IT with severe bleeding history and platelet ATP secretion defects. Importantly, SLFN14 expression is dramatically reduced in patients’ platelets (up to 80%) compared to healthy controls and in transfected cells (up to 95%), suggesting a dominant negative effect of mutants on the synthesis or stability of the wild-type (WT) form [Fletcher et al. 2015]. Shortly after this seminal study, Marconi et al. reported the fourth heterozygous missense mutation in *SLFN14* associated with IT [Marconi et al. 2016]. Notably, the affected residue R223W in AAA domain of SLFN14 is located nearby to previously reported mutations. This novel mutation was shown to mediate reduced proplatelet formation and decreased megakaryocyte maturation in patient derived megakaryocytes. Consistently with above mentioned data, SLFN14 expression was below 50% despite the heterozygous nature of the mutation [Marconi et al. 2016]. This finding supports the idea of dominant negative effect of mutant forms on the SLFN14 WT expression [Fletcher et al. 2015; Marconi et al. 2016]. However, due to the limited knowledge of the function of SLFN14, more detailed characterization of *SLFN14* and its role in platelet biogenesis is critically important.

Alongside these genetic studies, Pisareva et al. demonstrated the endoribonucleolytic activity of purified SLFN14 in biochemical experiments and suggested the role of protein in translation control [Pisareva et al. 2015]. More specifically, it was shown that SLFN14 associates with ribosomes and ribosomal subunits, and cleaves RNA, but preferably rRNA and ribosome-bound mRNA, in a Mg^2+^-dependent and NTP-independent manner. This leads to the degradation of ribosomal subunits [Pisareva et al. 2015]. More recently a more global approach [Mills et al. 2016] described a study of a ribosomal rescue pathway which involves both erythroid cells and platelets and the many proteins involved in this process such as proteins like SLFN14.

Based on the presence of characteristic slfn signature motifs, SLFN14 belongs to the Schlafen protein family, limited to mammals and encoded by six *SLFN* genes in humans [Geserick et al. 2004; Brady et al. 2005]. All family members comprise a conserved N-terminus containing a putative AAA domain implicated in ATP binding, but only longer forms of *SLFN* genes (including *SLFN14*) possess a C-terminal extension with motifs, which are specific for superfamily I DNA/RNA helicases [Geserick et al. 2004]. SLFN proteins are involved in T-cell development [Geserick et al. 2004; Schwarz et al. 1998; Berger et al. 2010], differentiation [Patel et al. 2009], and immune response [Li et al. 2012], but their exact cellular functions still remain elusive.

To get insights into the fundamental role of SLFN14, we aimed to assay endoribonuclease activity, intracellular distribution, and stability of WT and IT-related missense mutation forms of protein in human cells. We found that SLFN14 co-localizes with ribosomes, causes the endoribonucleolytic degradation of rRNA, and mediates total RNA clearance in cells. Mutations do not affect all tested activities of the protein, but dramatically reduce mutants’ stability at post-translational level and downregulate the co-expression of WT form. In light of our data, implications for the fundamental role of SLFN14 are discussed.

## RESULTS

### SLFN14 WT and IT-related missense mutation variants reveal the same subcellular distribution, co-localize with 5.8S rRNA, and cause ribosome degradation

To date, data on SLFN14 activity are limited by describing the protein as an endoribonuclease in a rabbit reticulocyte lysate and in a reconstituted *in vitro* mammalian translation system [Pisareva et al. 2015]. To expand our knowledge on the function of this protein, we aimed to analyze SLFN14 in transfected cell lines. Therefore, we used myc-tagged human SLFN14 WT (SLFN14(WT)-myc) and three previously reported missense mutation variants K218E, K219N, and V220D (SLFN14(K218E)-myc, SLFN14(K219N)-myc, and SLFN14(V220D)-myc,respectively) that were cloned into the mammalian expression vector for transient expression [Fletcher et al. 2015]. Dami cells, a human megakaryocytic leukemia cell line, were selected as a host cell line. Dami cells were differentiated into megakaryocyte like cells using Phorbol 12-myristate 13-acetate (PMA) then transiently transfected with SLFN14 (WT/mut)-myc constructs for 48 hours and were subsequently analysed by immunostaining. Expression of SLFN14 (WT)-myc revealed a diffuse cytoplasmic and nuclear localization with some punctate structures observed (Figure 1A). None of the mutations affected the subcellular distribution and staining pattern of the protein (Figures 1A and B). Such a pattern may indicate the co-localization of SLFN14 with non-uniform dispersed components of the cell. It was previously shown that SLFN14 strongly binds to purified ribosomes in the sucrose density gradient (SDG) centrifugation experiment [Pisareva et al. 2015]. Therefore, we suggested that ribosomes could be a binding partner for SLFN14 in the cell. Immunostaining experiments with the antibodies against 5.8S rRNA revealed that SLFN14 (WT)-myc significantly co-localizes with ribosomes, and the presence of mutations do not influence this co-localization (Figures 1A-C). SLFN14 was reported to possess the ribosomal binding site within 179 amino acids in the central part of the protein beside the AAA domain to the C-terminus [Pisareva et al. 2015]. Therefore we proposed that SLFN14 should bind directly to the ribosomes in cells. Moreover, ribosomal binding of SLFN14 indicates the role of protein in the translation control.

**Figure 1.**
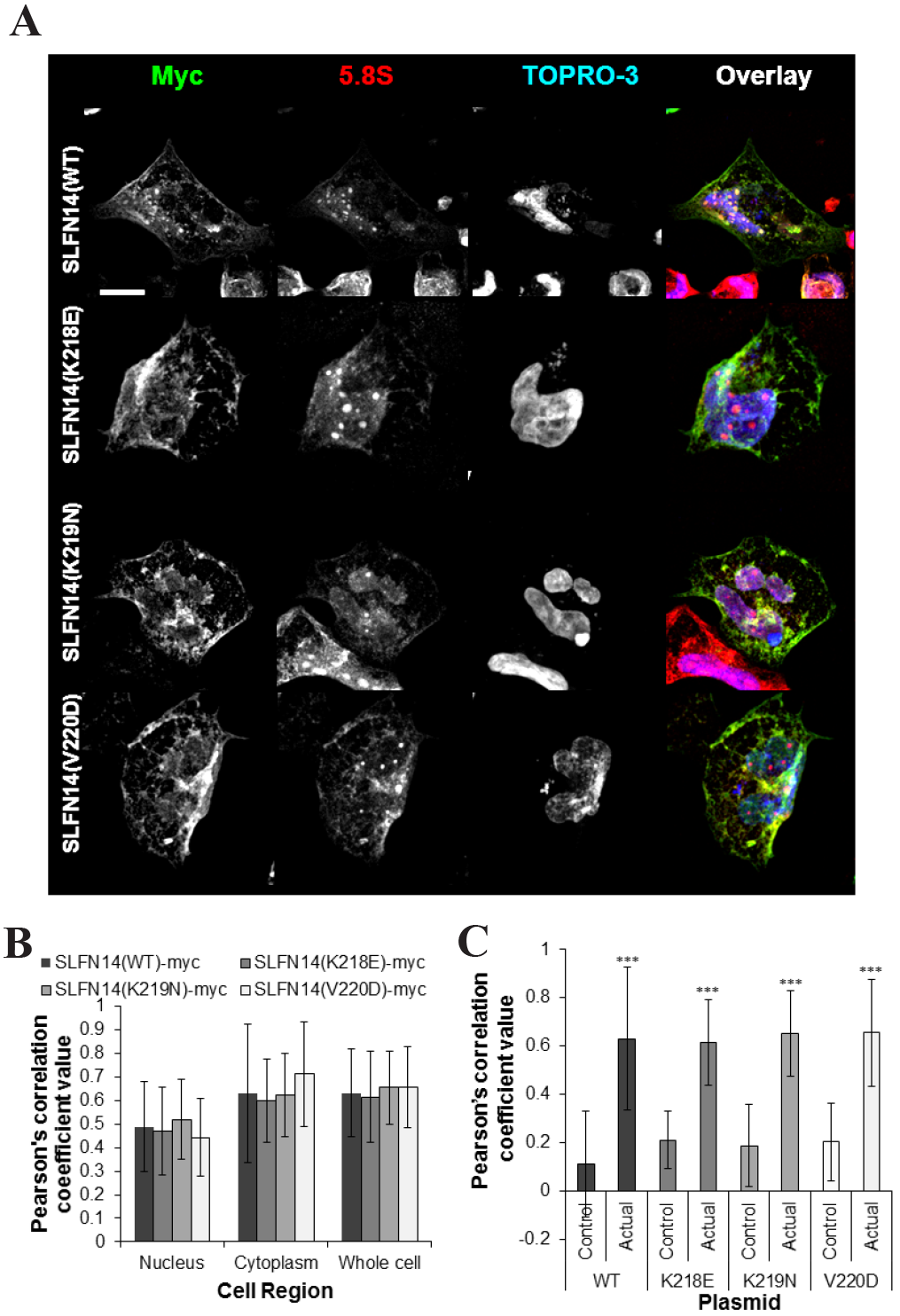
Significant co-localisation is observed between SLFN14 (WT/mut)-myc and 5.8S rRNA in differentiated Dami cells. There is no alteration in subcellular distribution between SLFN14 (WT)-myc and SLFN14 (mut)-myc distribution or co-localisation with 5.8S rRNA. (**A**) Transiently transfected differentiated Dami cells expressing SLFN14(WT/mut)-myc for 48 hours were probed with rabbit anti-myc and mouse anti-5.8S primary antibodies followed by incubation with anti-rabbit AlexaFluor488, anti-mouse AlexaFluor568 secondary antibodies and TO-PRO-3 Iodide nuclear stain. A representative image from 3 independent experiments, scale bar denotes 15μm. (**B**)Pearson’s correlation coefficient data demonstrating no change in co-localisation or subcellular-distribution between 5.8S rRNA and SLFN14 (WT)-myc or SLFN14 (mut)-myc in comparison to control areas. n=at least 40 cells analysed from 3 independent experiments. (**C**) Pearson’s correlation coefficient data demonstrating a significant increase in co-localisation between 5.8S rRNA and SLFN14 (WT/mut)-myc staining in comparison to control areas. *** P≤0.001 co-localisation between 5.8S rRNA and SLFN14 (WT/mut)-myc and control area. Error bars ± standard deviation.

Rabbit SLFN14 was previously shown to cause rRNA cleavage and ribosome degradation in a rabbit reticulocyte lysate and in a reconstituted *in vitro* mammalian translation system [Pisareva et al. 2015]. Notably, SLFN14 is highly homologous among all mammalian species. To test whether SLFN14 could provide ribosome degradation in the intact cells of human origin, we estimated 5.8S rRNA content in transiently transfected Dami cells expressing SLFN14 (WT/mut)-myc for 48 hours by immunostaining of 5.8S rRNA. For all SLFN14 variants, we detected about 50% to 70% statistically significant reduction of 5.8S rRNA content (Figure 2). Based on staining intensity values (Figure 2B), we cannot state that mutations compromise the ribosome degradation activity of SLFN14. However, taking into account of previous biochemical data [Pisareva et al. 2015], we suggest the more direct rather than auxiliary role of SLFN14 in the ribosome degradation.

**Figure 2.**
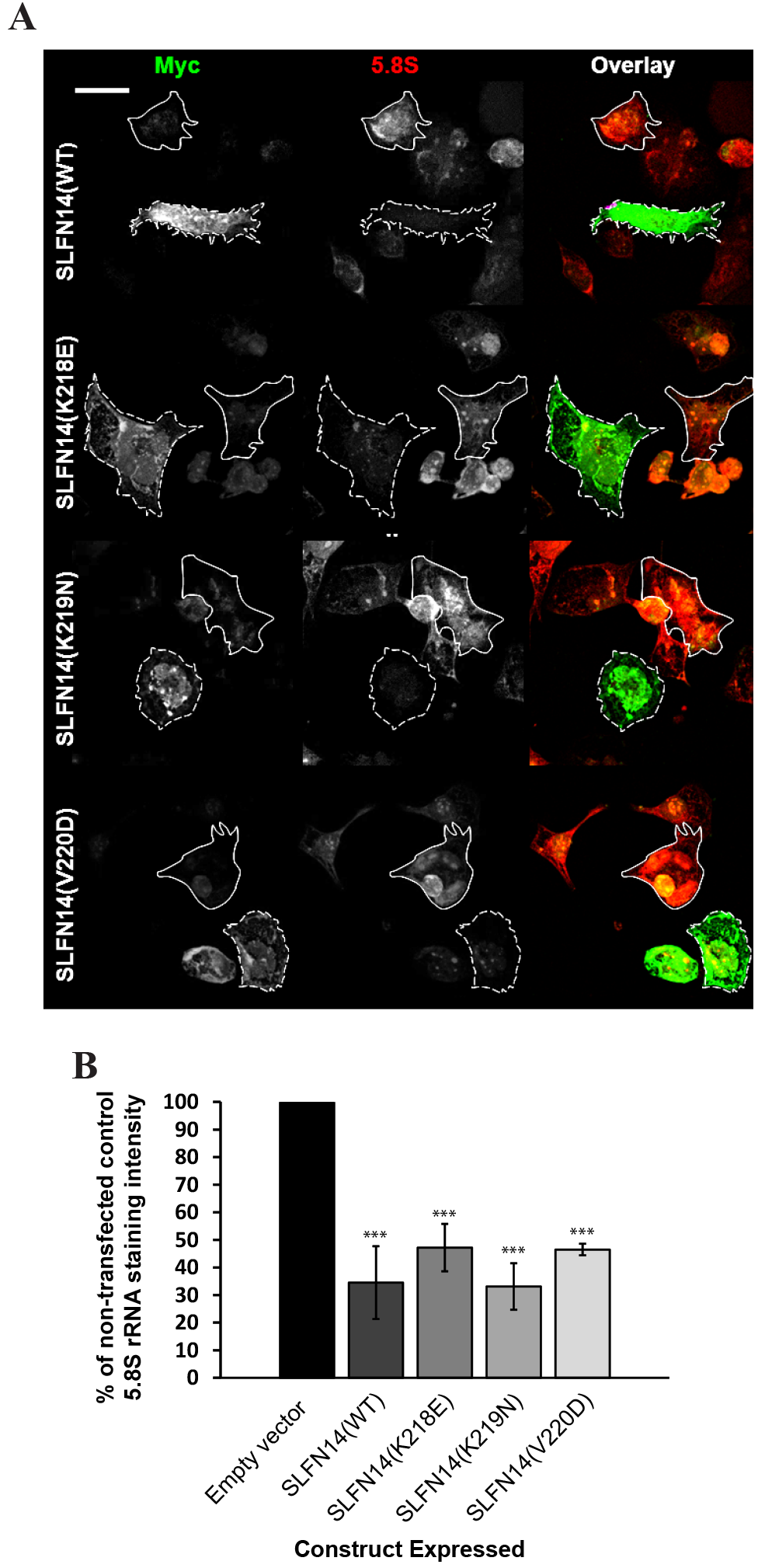
Reduced intensity of 5.8S rRNA staining in differentiated Dami cells expressing wild type/mutant SLFN14 constructs. (**A**) Transiently transfected Dami cells expressing SLFN14(WT/mut)-myc for 48 hours were probed with rabbit anti-myc and mouse anti-5.8S primary antibodies followed by incubation with anti-rabbit AlexaFluor488 anti-mouse AlexaFluor568 secondary antibodies. The dashed white line outlines cells expressing SLFN14 (WT/mut)-myc, the solid line represents the outline of cells which were nottransfected. A representative image from 3 independent experiments, scale bar denotes 15μm. (**B**) Average intensity measurements from the entire cell area were quantified from images represented in A.n=at least 40 cells analysed from 3 independent experiments. *** P≤0.001 when compared to non-transfected cells. Error bars ± standard deviation.

In conclusion, SLFN14 binds to the ribosomes mediating ribosome clearance in megakaryocyte-like cells, and IT-related missense mutations do not compromise these cellular activities of protein assuming that SLFN14 dysfunction takes place at a different molecular level.

### Overexpressed SLFN14 WT and mutants associate with ribosomes and individual ribosomal subunits causing the endoribonucleolytic cleavage of rRNA

To exclude that the discovered SLFN14 activities are related to a megakaryocyte-specific cell line, we also assayed the protein in HEK293T cells, a human embryonic kidney cell line. Based on reported and our current data, SLFN14 is suggested to be involved in translation control. A “gold standard” assay to evaluate the role ofa particular protein in translation control is to test its influence on the ribosomal profile. Therefore, transiently transfected HEK293T cells expressing SLFN14(WT/mut)-myc or harbouring empty vector (EV) for 48 hours were collected and lysed, and cell lysates were subjected to centrifugation through 10%-50% SDG in order to obtain a ribosomal profile (Figure 3A). Importantly, cells were lysed in the presence of cycloheximide to exclude polysome runoff during subsequent manipulations with the lysate. This antibiotic interferes with the translocation step during protein synthesis and, in such a way, blocks the translation elongation stage. Assignment of peaks in the ribosomal profile was made according to immunoblotting analysis of corresponding fractions by antibodies against large ribosomal protein RPL3 and small ribosomal protein RPS19 as exemplified by EV-transfected sample (Figure 3B).

**Figure 3.**
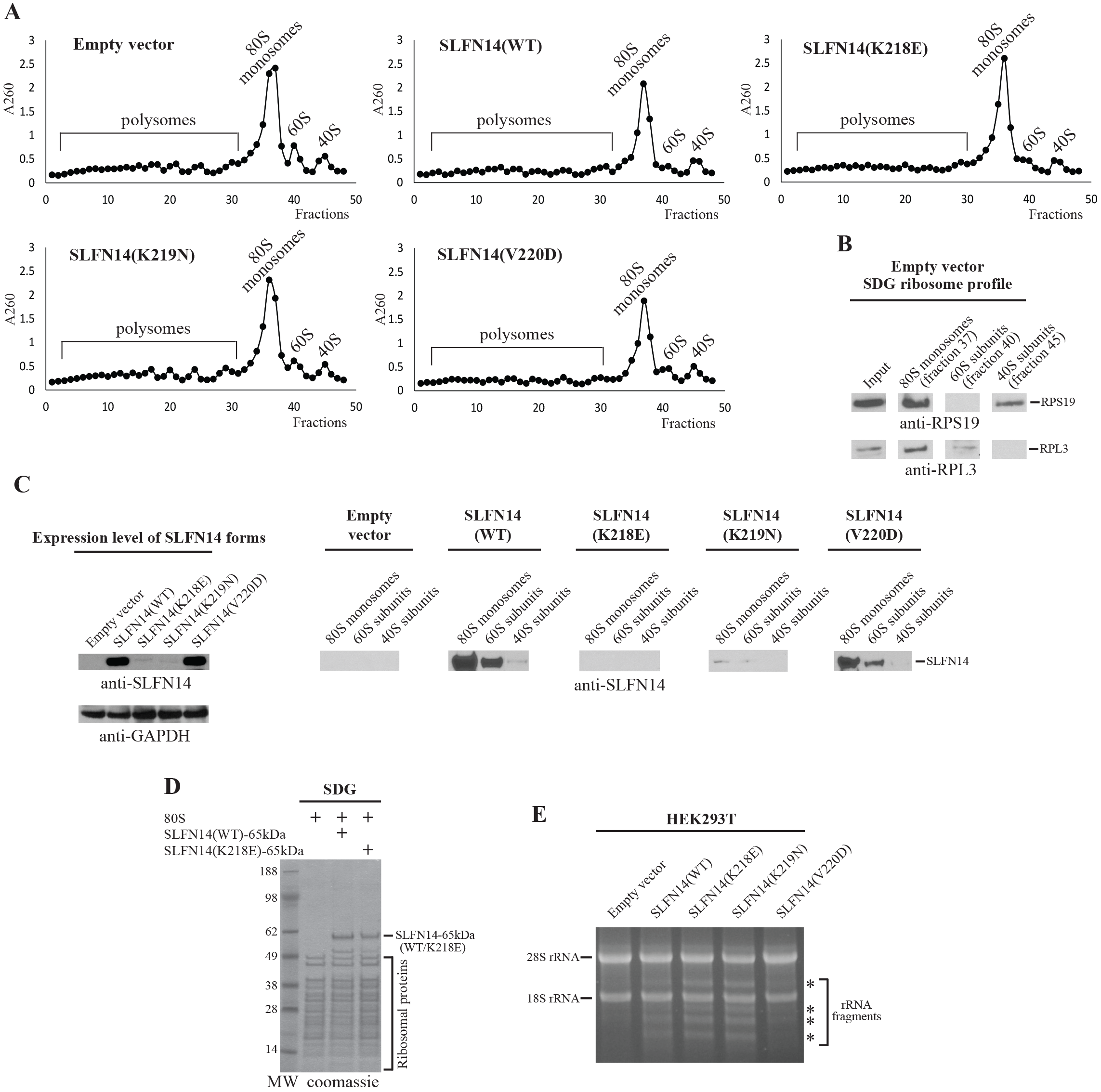
Association of SLFN14 (WT/mut)-myc with ribosomes and ribosomal subunits resulting in rRNA endonucleolytic degradation in HEK293T cells. (**A**) Ribosomal profiles of HEK293T cell lysates with overexpressed one of SLFN14 (WT/mut)-myc forms obtained by SDG centrifugation. (**B**) Assignment of ribosomal peaks in the ribosomal profile of empty vector transfected HEK293T cell lysate based on immunoblotting with anti-RPS19 andanti-RPL3 antibodies. (**C**) Expression levels of SLFN14 (WT/mut)-myc in HEK293T cells assayed by immunoblotting with anti-SLFN14 and anti-GAPDH (control) antibodies. Association of different forms of SLFN14 with 80S monosomes, 60S and 40S ribosomal subunits inthe corresponding fractions of ribosomal profiles (as depicted in A). Ribosomal fractions were concentrated and analyzed by immunoblotting with anti-SLFN14 antibodies. (**D**) Binding of recombinant SLFN14 (WT)-65kDa and SLFN14 (K218E)-65kDa to assembled 80S ribosomes assayed by SDG centrifugation and Coomassie staining. (**E**) rRNA degradation in SLFN14(WT/mut)-overexpressed HEK293T cells assayed by denaturing agarose/formaldehyde gel electrophoresis (n=3 independent experiments). The asterisks indicate the main bands of rRNA fragments. Positions of the 28S rRNA and 18S rRNA are shown.

Separation of ribosomal fractions in the ribosomal profile permitted us to assay the affinity of SLFN14 to 80S ribosomes and individual ribosomal subunits. All SLFN14 forms except SLFN14 (K218E)-myc bind predominantly to 80S ribosomes and 60S ribosomal subunits (Figure 3C). The weak signals of SLFN14 (K219N)-myc and the absence of SLFN14 (K218E)-myc in ribosomal peaks correlate with the low expression level of proteins (Figure 3C).It was previously shown that the expression level of SLFN14 (K218E)-myc in HEK293T cells is the lowest among all the described mutants [Fletcher et al. 2015]. Therefore, we hypothesized that the absence of SLFN14 (K218E)-myc in ribosomal fractions is a result of the low content and/or continuous degradation during cell lysate manipulation rather than of the compromised ribosomal binding activity. To test our hypothesis, we utilized a previously described E.coli expression vector for a 65kDa C-terminally truncated His-tagged form of human SLFN14 (SLFN14-65kDa) [Pisareva et al. 2015]. This was the longest form of SLFN14, which was available in a soluble state after expression, whereas all longer forms completely precipitated [Pisareva et al. 2015]. We introduced the corresponding mutation into SLFN14-65kDa to obtain K218E mutant expression vector (SLFN14 (K218E)-65kDa). Due to limited solubility, SLFN14-65kDa or SLFN14 (K218E)-65kDa proteins in the form of eluates after Ni-NTA resin were mixed with 80S ribosomes reconstituted from purified 40S and 60S ribosomal subunits. The reaction mixture was subjected to the centrifugation through SDG to separate 80S ribosomes from unbound components, and the 80S ribosomal peak was assayed by denatured PAGE and Coomassie staining. Both SLFN14-65kDa and SLFN14 (K218E)-65kDa were found associated with 80S ribosomes (Figure 3D). Therefore, none of the SLFN14 mutations affect the ribosomal binding activity of SLFN14 in different cell lines.

In the immunostaining experiment, all SLFN14 forms reduced the intensity of 5.8S rRNA staining in Dami transfected cells that could be a result of ribosome degradation. In our next experiment, we raised several key questions. Could the rRNA degradation process be involved in the elimination of ribosomes? Is the degradation activity of SLFN14 cell type-specific? How does the rRNA pattern look after the overexpression of SLFN14? To answer these questions, transiently transfected HEK293T cells expressing SLFN14(WT/mut)-myc or harbouring EV for 48 hours were collected, total RNA was isolated, and the equal amounts of total RNA from each sample were assayed by denaturing agarose/formaldehyde gel electrophoresis. In contrast to EV, overexpression of any SLFN14 form resulted in the characteristic pattern of rRNA endoribonuclealytic degradation (Figure 3E). In summary, the endoribonucleolytic activity of SLFN14 should relate to the observed ribosome degradation in transfected cells. Notably, IT-related missense mutations do not affect rRNA cleavage pattern pointing to that SLFN14 dysfunction in platelet biogenesis cannot be explained by the compromised endoribonucleolytic activity of protein.

### *SLFN14* missense mutations cause the low expression of mutants as a result of post-translational degradation due to partial misfolding, and implicate SLFN14 WT into the degradation through the formation of oligomeric forms

One of the interesting reported findings is that SLFN14-related IT patients displayed 65%-80% reduction in SLFN14 protein level despite the heterozygosity, assuming that the mutant allele influences the synthesis and/or stability of both mutant and WT proteins [Fletcher et al. 2015; Marconi et al. 2016]. This effect was confirmed in overexpression studies in transfected cells [Fletcher et al. 2015]. Therefore, in our next set of experiments, we aimed to determine the stage, at which mutant expression could be affected, and to understand how mutants could influence the SLFN14 WT protein level. For that, we employed above mentioned SLFN14 (WT/mut)-myc vectors and a newly constructed SLFN14 (WT)-GFP mammalian expression vector containing the human SLFN14 coding region with a GFP tag. Transiently transfected HEK293T cells co-expressing SLFN14 (WT)-GFP and either the SLFN14 (WT/mut)-myc vectors for 48 hours were analysed by immunoblotting. First, consistently with published data [Fletcher et al. 2015], we found that SLFN14 (K218E)-myc, SLFN14 (K219N)-myc, and SLFN14 (V220D)-myc expression was reduced to 10%, 10%, and 54% of SLFN14(WT)-myc expression, respectively (Figure 4A and B). Second, SLFN14(WT)-GFP protein level dropped to 37% and 54% upon co-expression of that with SLFN14(K218E)-myc and SLFN14(K219N)-myc, respectively, compared to co-expression with SLFN14(WT)-myc control (Figure 4A and B). These data correlate with the proposed dominant negative effect of SLFN14 mutants. The detected elevation of SLFN14 (WT)-GFP protein level upon co-expression with SLFN14 (V220D)-myc compared to that with the SLFN14 (WT)-myc control was not statistically significant and we therefore suggest that this mutant behaves in a different way (Figure 4A and B).

**Figure 4.**
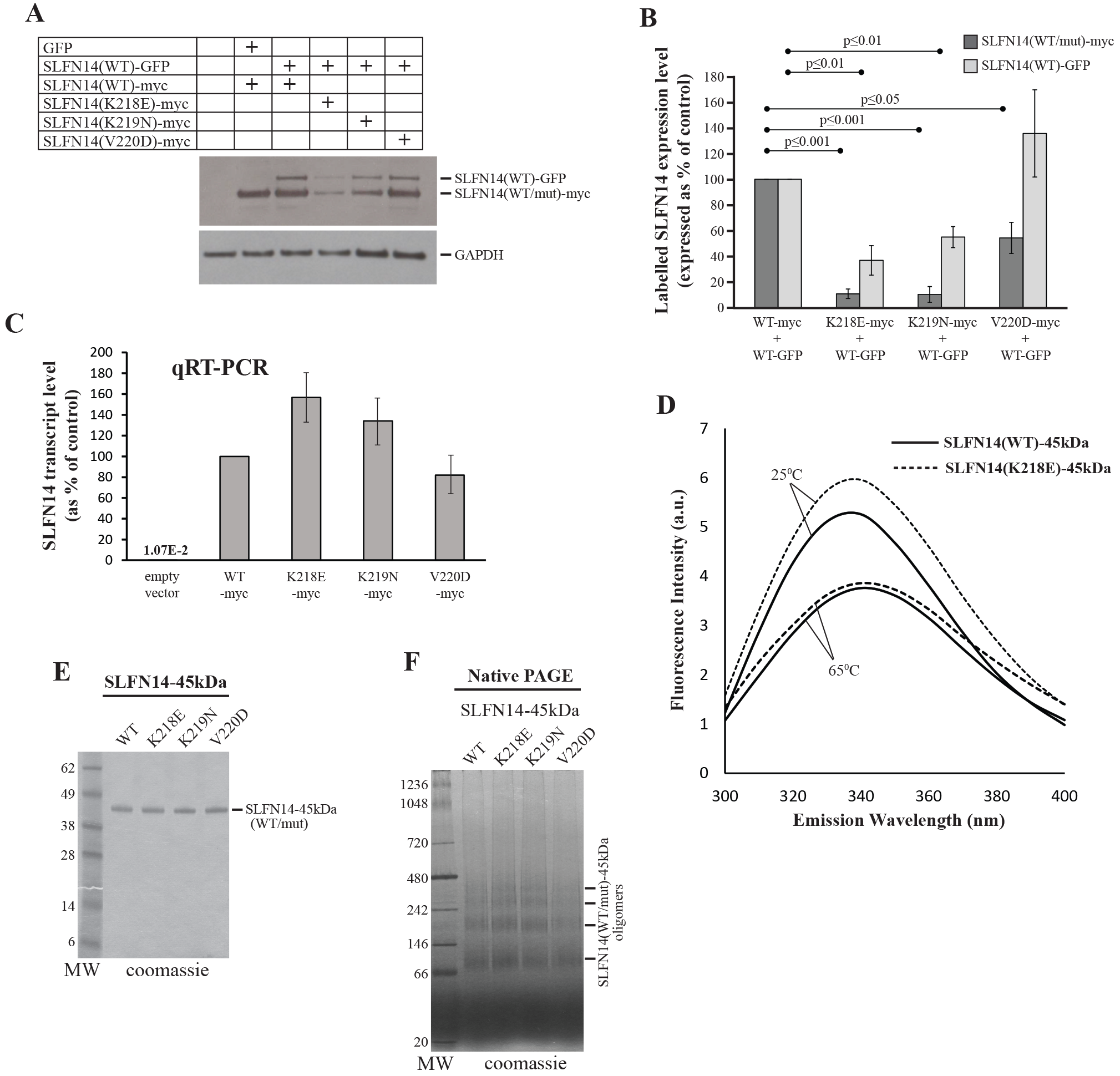
Role of *SLFN14* missense mutations in protein expression. (**A**) Immunoblotting image showing levels of SLFN14 (WT/mut)-myc and SLFN14(WT)-GFP in HEK293T cells transiently expressing the above constructs. The blot was probed with anti-SLFN14 and anti-GAPDH primary antibodies followed by incubation with anti-rabbit HRP. (**B**) Quantification of SLFN14 (WT/mut)-myc and SLFN14 (WT)-GFP protein expression from immunoblotting analysis of n=3 lysate samples percondition from 3 independent experiments. All values are mean ± standard deviation. (**C**) Quantification of relative SLFN14 (WT/mut)-myc transcript level normalized to GAPDH control transcript from qRT-PCR test data on n=6 independent sets of lysate samples. All values are mean ± standard deviation. (**D**) Fluorescence emission spectra of SLFN14 (WT)-45kDa and SLFN14 (K218E)-45kDa proteins collected at the same protein concentration at two different temperatures: 25°C and 65°C (n=3 independent experiments). (**E**) Purifiedrecombinant SLFN14-45kDa WT and mutants resolved by SDS-PAGE. (**F**) Oligomerization capacity of SLFN14-45kDa WT and mutants assayed by native PAGE. Positions ofdifferent oligomeric forms are shown.

The regulation of expression of SLFN14 mutants should take place at either transcriptional or post-translational level. To evaluate the effect of mutations on the RNA level, we employed qRT-PCR. We isolated total RNA from HEK293T cells expressing one of SLFN14 (WT/mut)-myc variants, and assayed samples for expression of *SLFN14* and *GAPDH* control transcripts. The difference in a normalized transcript concentration between *SLFN14* WT and each mutant form did not exceed 1.6 fold (Figure 4C). Importantly, the endogenous *SLFN14* transcript was not detected (Figure 4C). Taking into account a strong reduction of SLFN14 (K218E)-myc and SLFN14 (K219N)-myc expression, we conclude that the regulation of expression should occur at a post-translational rather than a transcriptional stage, at least for these two mutants. We then hypothesized that missense mutations could affect protein folding. To test this hypothesis, we compared folding between the SLFN14 WT and K218E form, which revealed the most dramatic effect on its own expression and WT form co-expression in our experiments. Tertiary structure of protein can be analyzed using a fluorescence spectroscopy technique based on intrinsic protein fluorescence. Two amino acids, Trp and Tyr, are experimentally used to obtain a strong fluorescent signal. The emission energy of these residues is highly sensitive to the polarity of the environment. In the native folded conformation, Trp and Tyr are generally hidden in the hydrophobic core of the protein giving high intensity fluorescence signal. In contrast, a hydrophilic environment results in a low intensity fluorescence signal. The protein sample is excited at 280 nm wavelength, and fluorescence spectrum is collected in a 300-400 nmwavelength range. For the experiment, we employed the already described E.coli expression vector for 45kDa C-terminally truncated His-tagged form of human SLFN14 (SLFN14-45kDa) [Pisareva et al. 2015]. Compared to SLFN14-65kDa and longer protein forms, SLFN14-45kDa could be purified in a large amount with high homogeneity [Pisareva et al. 2015]. We inserted the corresponding mutation into SLFN14-65kDa to obtain K218E mutant expression vector (SLFN14 (K218E)-45kDa). Analysis of SLFN14-45kDa and SLFN14(K218E)-45kDa fluorescence spectra revealed the difference in the maximum of emission energy at 340 nm wavelength indicating different conformations of proteins and confirming our hypothesis(Figure 4D). In the control experiment, after incubation of proteins at 65°C resulting in their denaturation, maximum intensities of emission matched (Figure 4D). In conclusion, we state that *SLFN14* missense mutations lead to post-translational degradation of mutants as a result of partial protein misfolding.

Taking into account a post-translational regulation of *SLFN14* expression, we suggested that mutants could involve a WT protein into degradation through the formation of heterogeneous oligomeric forms. Indeed, AAA proteins form oligomeric assemblies, mostly homo-hexamers, which are critical for their activities [Ogura and Wilkinson 2001]. To evaluate the SLFN14 tendency to oligomerization, we assayed the mobility of protein in native PAGE. We used the above mentioned SLFN14-45kDa and SLFN14 (K218E)-45kDa proteins as well as newly constructed, E.coli expressed and purified SLFN14 (K219N)-45kDa and SLFN14 (V220D)-45kDa mutants (Figure 4E). As a result, SLFN14 WT migrated in a native PAGE in the form of homo-oligomers of different orders, and none of mutations affected the protein pattern (Figure 4F). Therefore, we suggest that our findings underlie the mechanism of *SLFN14* expression regulation in transfected cells.

## DISCUSSION

*SLFN14* is one of the most recently discovered genes known to cause inherited thrombocytopenia. Four missense mutations of *SLFN14* are identified to date, linked to dysregulated platelet maturation and platelet dysfunction, resulting in disproportionate bleeding in affected patients. Schlafen family members are poorly studied and their functions are not completely understood making the cellular role of SLFN14 hard to predict. The only functional characterization report describes SLFN14 as an endoribonuclease in a rabbit reticulocyte lysate. To advance our knowledge on SLFN14, we characterized the protein in different transfected cell lines.

Consistent with published data, we detected a diffuse immunostaining pattern for overexpressed SLFN14 WT throughout the nucleus and cytoplasm with some punctate structures in Dami cells. Immunostaining assay also revealed that the non-uniform distribution of SLFN14 WT is a result of co-localization with 5.8S rRNA indicating the ribosome as a binding partner for the protein in the cell. This finding unambiguously points to the role of SLFN14 in translation control. Taking into account the reported biochemical data, we assayed the rRNA degradation capacity of protein in question, and found that SLFN14 WT overexpression leads to rRNA endoribonucleolytic cleavage and degradation in Dami and HEK293T cells, which represent megakaryocyte-related and unrelated cell lines, respectively. Therefore, SLFN14 is a bona fide mammalian endoribonuclease, and the endoribonucleolytic activity of protein does not depend on cell-specific co-factors. Importantly, only few endoribonucleases, involved in the translation control, have been described so far. That is because it is hard to predict the endoribonucleolytic activity of protein based on its primary sequence due to a high variety in the organization of the active center and, thus, a wide structural diversity of this class of enzymes. Notably, IT-related missense mutations K218E, K219N, and V220D do not affect the distribution, 5.8S rRNA co-localization, endoribonucleolytic activity, and ribosome clearance-mediated function of SLFN14 in the cell. This data implies that protein dysfunction in platelet biogenesis takes place at a different cellular level.

Analysis of the ribosomal profiles revealed that all SLFN14 forms bind to ribosomes. Notably, as follows from the experiment in the binary system with the recombinant protein, ribosomal association of SLFN14 is direct and not mediated by co-factors. This is the first reported case of direct ribosomal association of endoribonuclease within the cell.

It was reported that *SLFN14* missense mutations cause the decreased protein expression in transfected cells and downregulate the expression of SLFN14 WT form in patients. Consistently, in our study, the protein level dropped dramatically for K218E and K219N mutants and moderately for V220D mutant in HEK293T cells. Moreover, GFP-tagged SLFN14 (WT) protein levels reduced by 3 and 2 times upon co-expression with K218E and K219N mutants, respectively, compared to co-expression with WT control. qRT-PCR data on the SLFN14 transcript levels displayed that regulation of expression takes place at a post-translational rather than a transcriptional stage. We hypothesized that the missense mutations could cause partial misfolding of protein. Indeed, to preven tthe potentially hazardous effect, the cell employs several degradation pathways to destroy improperly folded proteins [Nedelsky et al. 2008; Smith et al. 2011; Varshavsky 2012]. Interestingly, protein misfolding is shown to underlie hundreds of diseases [Valastyan and Lindquist 2014]. Fluorescence spectroscopy demonstrated different tertiary structures of SLFN14 WT and K218E supporting our suggestion that post-translational degradation of mutants is a result of partial protein misfolding. But how do SLFN14 mutants mediate SLFN14 WT degradation? All AAA proteins, which SLFN14 belongs to, tend to form oligomers in the cell. Consistently, native PAGE revealed that SLFN14 WT forms oligomers of different order, and mutations do not affect the protein pattern in the gel. Therefore, SLFN14 mutants could cause increased degradation of wild type SLFN14 by forming hetero-oligomers of wild type/mutant SLFN14 leading to instability of the entire protein complex, and could explain a dominant-negative effect of mutant allele on SLFN14 WT expression in patients.

What are the implications of our findings for platelet biogenesis and IT-related dysregulation of this process? It is well known that mature platelet and erythrocytes have only residual “RNA content” levels, if any [Hamilton 2010; Angénieux et al. 2016]. Here we show that SLFN14 reveals the endoribonucleolytic activity resulting in rRNA cleavage and degradation in different transfected cells. Therefore, we cautiously speculate that SLFN14 causes RNA degradation and, in such a way, mediates RNA clearance during platelet and erythrocyte maturation. Interestingly, SLFN14 was found abundant in rabbit reticulocytes, but below the detection limit in rabbit liver, lung and brain tissues [Pisareva et al. 2015]. Moreover, endogenous SLFN14 expression was shown to be undetectable or extremely low in HEK293, HEK293T, HeLa, CEM, and Jurkat cells [Li et al. 2012]. On the other hand, in the independent studies, SLFN14 was demonstrated to be involved in platelet biogenesis. Taking these data together, we again cautiously hypothesize that SLFN14 is specifically expressed and acts during these two blood cell types maturation processes.

Interestingly a recently published and more global approach [Mills et al. 2016] was studied to outline a ribosomal rescue pathway which involves both erythroid cells and platelets and the many proteins involved in this process such as proteins not dissimilar to SLFN14. Therefore, our more specific study follows on from this and complements this study by implicating a ribosomal pathway more specifically in SLFN14 as outlined here.

Since we did not detect any difference between SLFN14 WT and mutants in our experiments, except at the expression level, we think that the dysregulation of thrombopoiesis is a result of improper degradation of mutants. There are examples of human genetic diseases caused by the degradation of mutant proteins despite these proteins retaining their functionality [Valastyan and Lindquist 2014]. A canonical example is the disease cystic fibrosis linked to a single phenylalanine residue deletion at position 508 of cystic fibrosis transmembrane conductance regulator protein targeting a misfolded protein for degradation [Qu et al. 1997]. Another case of improper degradation-associated disease is a Gaucher’s disease, which is caused by a variety mutations in β-glucosidase [Futerman and van Meer G. 2004; Cox and Cachón-González 2012]. In conclusion, our findings contribute to the understanding of the mechanism underlying platelet biogenesis in general and SLFN14-related inherited thrombocytopenia more specifically.

## MATERIALS AND METHODS

### Plasmids

Mammalian expression vector containing the full coding region of human *SLFN14* and GFP-tag was purchased from GeneCopoeia (SLFN14(WT)-GFP). Mammalian expression vector for human SLFN14(WT)-myc as well as *E.coli*-based expression vectors for His-tagged C-terminal deletion mutants of human *SLFN14* (SLFN14(WT)-65kDa and SLFN14(WT)-45kDa) have been previously described [Pisareva et al. 2015; Fletcher et al. 2015]. The *SLFN14* missense mutations K218E, K219N and V220D were created by site-directed mutagenesis of corresponding vectors.

### Antibodies

We used myc (Cell Signaling Technology, #9B11), myc (Abcam, #9106), 5.8S rRNA (Abcam, #ab37144) SLFN14 (Abcam, #ab106406), RPS19 (Bethyl Laboratories, #A304-002A), RPL3(Bethyl Laboratories, #A305-007A), GFP (Sigma, #G1544), GAPDH (Abcam, #ab9485), anti-rabbit AlexoFluor488 (ThermoFisher Scientific, #A-11034) and anti-mouse AlexoFluor568 (ThermoFisher Scientific, #A-11004) antibodies.

### Cell culture, plating and transfection

HEK293T cells were cultured in DMEM plus L-glutamine (Invitrogen) (plus 10% fetal calf serum, 1% pen/strep (both from GIBCO)). Dami cells were cultured in RPMI (GIBCO) plus 1% L-glutamine, 10% fetal calf serum and 1% pen/strep. Cells were plated into 6 well plates with/without sterilized 23mm glass coverslips at a density of 5×10^5^ cells/ml. Dami media was supplemented with PMA (Phorbol 12-myristate 13-acetate (Sigma-Aldrich)) to a final concentration of 10μM. Cells were transfected 24 hours post-plating with 5.5μl (1mg/ml pH 7.4 PEI (Polyethyleneimine (Sigma-Aldrich)), 1.2μg DNA and 140μl Optimem (Invitrogen) per well of a 6 well plate and used in studies 48 hours post-transfection.

### Purification of ribosomal subunits

Native rabbit 40S and 60S subunits were purified as described [Pisarev et al. 2007].

### Purification of *E.coli*-expressed SLFN14 WT and mutants

Recombinant SLFN14(WT/mut)-45kDa were expressed in 1 L of *E. coli* BL21(DE3) media after induction with 0.1 mM IPTG for 16 h at 16°C. After expression, the proteins were isolated by affinity chromatography on Ni-NTA agarose followed by FPLC on a MonoS column. FPLC fractions were collected across a 100-500 mM KCl gradient. SLFN14(WT/mut)-45kDa were eluted in the range of 210-250 mM KCl.

Recombinant SLFN14(WT/K218E)-65kDa were expressed in 1 L of *E. coli* BL21(DE3) media after induction by 0.1 mM IPTG for 16 h at 16°C and purified on Ni-NTA agarose according to manufacturer’s protocol.

### HEK293T cell extract preparation

To prepare cell extract, transiently transfected HEK293T cells expressing SLFN14(WT/mut)-myc or harbouring empty vector were cultured, plated and transfected as described previously. Prior to use cells were washed 3 times with 1 × PBS pH 7.4. 3 × 10^5^ HEK293T cells were resuspended in 300 μl of pre-chilled Tris-based lysis buffer (20 mM Tris-HCl pH 7.5, 100 mM potassium chloride, 2.5 mM magnesium chloride, 1 mM DTT, 0.1 mg/ml cycloheximide, 0.5 % Triton X-100, 40 units/ml DNAse I from NEB, HALT protease inhibitor cocktail EDTA-free from Thermo Scientific). The cells were allowed to swell for 10 minutes on ice and centrifuged at 16,000 g for 10 min at 4°C.

### Immunoblot analysis and densitometry

Transiently transfected HEK293T cells expressing SLFN14(WT/mut)-myc/myc empty vector and SLFN14(WT)-GFP were lysed as above and analysed using densitometry after immunoblotting. Western blot band intensity was quantified in NIS Elements version 4.00 .07 as follows: the ROI selection tool was used to draw around the largest band and the average intensity was measured. This box was used to measure the average band intensity of other bands. Background intensity was measured using the same ROI box moved to 4x non-band region in the same lane as the band measured. These values were logged to Excel, for both SLFN14 and GAPDH the average band value was then subtracted from the average background value. To correct for minor differences in protein levels seen in the GAPDH protein control, the band value for SLFN14 was divided by the average band value for GAPDH.

### HEK293T ribosomal profile preparation

To obtain the ribosomal profile, 100 μl HEK293T cell extract was diluted with 300 μl buffer A (20 mM Tris-HCl pH 7.5, 100 mM KCl, 2.5 mM MgCl_2_, 1 mM DTT, 0.1 mg/ml cycloheximide) and subjected to centrifugation through 10-50 % SDG prepared in buffer B (20 mM Tris-HCl pH 7.5, 100 mM KCl, 15 mM MgCl2, 1 mM DTT, 0.1 mg/ml cycloheximide) in a Beckman SW41 rotor at 35,000 rpm for 150 minutes at 4°C. After centrifugation, 200 μl fractions were collected. Ribosomal fractions were concentrated and analyzed by immunoblotting.

### Ribosomal binding assay

SLFN14(WT)-65kDa or SLFN14(K218E)-65kDa proteins in the form of eluates after Ni-NTA resin were mixed with 50 pmol 80S ribosomes, reconstituted from purified 40S and 60S ribosomal subunits, in the presence of 1 mM AMPPNP. The reaction mixture was subjected to centrifugation through 10-30% SDG prepared in buffer C (20 mM Tris-HCl pH 7.5, 100 mM KCl, 2.5 mM MgCl_2_, 1 mM DTT) in a Beckman SW55 rotor at 53,000 rpm for 75 min at 4°C. 80S ribosomal peak was assayed by NuPAGE 4-12% Bis-Tris SDS-PAGE (Invitrogen) and SimplyBlue SafeStain (Invitrogen) staining.

### rRNA degradation

To study rRNA degradation in HEK293T cell extract, total RNA was isolated with TRIzol reagent (Invitrogen) according to manufacturer’s protocol and 1 μg ofthat was analyzed by denaturing agarose/formaldehyde gel electrophoresis.

### Denaturing agarose/formaldehyde gel electrophoresis

RNA samples were loaded in a loading buffer (5x = 4 mM EDTA, 0.9 M formaldehyde, 20 % glycerol, 30.1 % formamide, 4x FA buffer, 0.4 μg/ml bromphenol blue) onto 1.2 % denaturing agarose/formaldehyde gel (3 % formaldehyde) prepared in FA buffer (20 mM MOPS, pH 7.0, 5 mM NaAc, 1 mM EDTA, 0.1 μg/ml ethidium bromide) and were resolved in FA buffer for 22 min at 4°C at 200V. After electrophoresis, gel was stained 2 times for 8 min each with 0.5 μg/ml ethidium bromide solution in water, washed 3 times for 5 min each with water and analyzed using shortwave UV (254 nm).

### qRT-PCR

To evaluate the relative SLFN14 transcript level in cells, 1 μg of isolated total RNA was converted into cDNA using random hexamers and SuperScript III kit (Invitrogen). 10 ng cDNA was subjected to real-time PCR employing iQ SYBR Green Supermix kit(Bio-Rad) with primers SLFN14 qPCR dir 5’-GCAAAGAAGTGGTTGGATGTAAG-3’, SLFN14 qPCR rev 5’-TCACAGCAGAAGTGGAATGTAG-3’, GAPDH qPCR dir 5’-GGTGTGAACCATGAGAAGTATGA-3’ GAPDH qPCR rev 5’-GAGTCCTTCCACGATACCAAAG-3’, and CFX96 Touch Real-Time PCR Detection System (Bio-Rad). SLFN14 transcription level was normalized to GAPDH control.

### SLFN14 oligomerization assay

5 μg SLFN14(WT/mut)-45kDa were analyzed by non-denaturing NativePAGE 4-16% Bis-Tris PAGE (Invitrogen) and SimplyBlue SafeStain (Invitrogen) staining in the presence of NativeMark Unstained Protein Standard (Invitrogen).

### Steady-state fluorescence measurements

Fluorescence emission spectra were obtained on a Fluoromax-3 spectrophotometer (Jobin Yvon Inc., Edison, NJ). 6 μg SLFN14(WT)-45kDa or SLFN14(K218E)-45kDa in a 200 μl buffer C were incubated at 25°C or 65°C for 10 min. Protein fluorescence was monitored using an excitation wavelength of 280 nm and an emission wavelength range 300-400 nm.

### Immunocytochemistry

Dami cells (American Type Culture Collection) were plated onto glass coverslips and transfected with SLFN14(WT/mutant)-myc as described above. Cells were fixed in 4% PFA for 5 minutes, and permeabilised in 0.1% Triton-X-100 in PBS for 5 minutes. Cells were incubated in block buffer (PBS (Invitrogen), 10% goat serum (Gibco), 5% BSA) for 1 hour. Cells were then incubated for 1 hour in primary antibody diluted in block as manufacturer’s instructions. Cells were washed in PBS and incubated for 1 hour in secondary antibody plus Topro-3 (Invitrogen) diluted as manufacturer’s instructions in block. Cells were mounted on glass slides using Hydromount mounting media (National Diagnostics).

### Immunofluorescence microscopy and analysis

All images were taken using a DM IRE2 Leica inverted microscope, SP2 confocal system running Leica Confocal Software Version 2.61 Build 1537. Confocal imaging was performed using the 488nm line of an Argon-Ion laser 457-514nM (to image AlexaFluor488 labelled constructs) and the 568 and 633 line of the HeNe lasers (to image AlexoFluor558 labelled constructs and TOPRO-3) with an HCX Plan Apo Ibd.BL 63x NA 1.4, Olympus objective. Z-stack images were taken at 10 slices per cell. Images were analyzed using NIS Elements Software.

Colocalization between SLFN14(WT/Mut)-myc and 5.8S rRNA was performed as follows:From average intensity projections the automated ROI tool in NIS-Elements, the entire cell volume, the cell cytoplasm and the nucleus of each cells was selected. Subsequently a Pearson’s coefficient comparing SLFN14(WT/Mut)-myc and 5.8S rRNA signals in these regions were obtained. In order to control for random colocalization, the 5.8S RNA image was rotated by 5°C and a Pearson’s coefficient repeated. All values were logged to Excel (Microsoft) A Student’s Test was performed to ascertain statistical significance between colocalization within the various subcellular regions for wild type of mutant SLFN14 and as a control between the rotated and non-rotated images.

Intensity measurements: Average intensity projections were created and ROIs drawn around the entire cell volume of a brightfield image of both untransfected cells and cells expressing SLFN14(WT/Mut)-myc images within the same field of view. This ROI outline was superimposed over the 5.8S RNA stained image and the average intensity measurement within the ROI calculated. All values were logged to Excel (Microsoft) A Student’s Test was performed to ascertain statistical significance between 5.8S RNA staining intensity between transfected and non-transfected cells.

## ACKNOWLEDGEMENTS

We thank the families for providing samples and our clinical and laboratory colleagues for their help. This work was supported by the National Institutes of Health (GM097014 to A.V.P.) and the British Heart Foundation (PG/13/36/30275 to N.V.M). The authors declare that they have no conflicts of interest with the contents of this article.

## REFERENCES

Nurden AT, Nurden P. 2007. Inherited thrombocytopenias. Haematologica 92:1158–1164.

Favier R, Raslova H. 2015 Progress in understanding the diagnosis and molecular genetics of macrothrombocytopenias. Br J Haematol 170:626–639.

Levin C, Koren A, Pretorius E, Rosenberg N, Shenkman B, Hauschner H, Zalman L, Khayat M, Salama I, Elpeleg O, et al. 2015 Deleterious mutation in the FYB gene is associated with congenital autosomal recessive small-platelet thrombocytopenia. J Thromb Haemost13:1285–1292.

Johnson B, Fletcher SJ, Morgan NV. 2016a. Inherited thrombocytopenia: novel insights into megakaryocyte maturation, proplatelet formation and platelet lifespan. Platelets 27:519–525.

Johnson B, Lowe GC, Futterer J, Lordkipanidzé M, MacDonald D, Simpson MA, Sanchez-Guiú I, Drake S, Bem D, Leo V, et al. 2016b. Whole exome sequencing identifies genetic variants in inherited thrombocytopenia with secondary qualitative function defects. Haematologica 101:1170–1179.

Pecci A. 2016 Diagnosis and treatment of inherited thrombocytopenias. Clin Genet 89:141–153.

Fletcher SJ, Johnson B, Lowe GC, Bem D, Drake S, Lordkipanidzé M, Guiú IS, Dawood B, Rivera J, Simpson MA, et al. 2015 SLFN14 mutations underlie thrombocytopenia with excessive bleeding and platelet secretion defects. J Clin Invest 125:3600–3605.

Marconi C, Di Buduo CA, Barozzi S, Palombo F, Pardini S, Zaninetti C, Pippucci T, Noris P, Balduini A, Seri M, et al. 2016 SLFN14-related thrombocytopenia: identification within a large series of patients with inherited thrombocytopenia. Thromb Haemost 115:1076–1079.

Pisareva VP, Muslimov IA, Tcherepanov A, Pisarev AV. 2015 Characterization of Novel Ribosome-Associated Endoribonuclease SLFN14 from Rabbit Reticulocytes. Biochemistry 54:3286–3301.

Mills EW, Wangen J, Green R, Ingolia NT. 2016 Dynamic Regulation of a Ribosome Rescue Pathway in Erythroid Cells and Platelets. Cell Rep 17:1–10.

Geserick P, Kaiser F, Klemm U, Kaufmann SHE, Zerrahn J. 2004 Modulation of T cell development and activation by novel members of the Schlafen (slfn) gene family harbouring an RNA helicase-like motif. Int Immunol 16:1535–1548.

Brady G, Boggan L, Bowie A, O’Neill LAJ. 2005 Schlafen-1 causes a cell cycle arrest by inhibiting induction of cyclin D1. J Biol Chem 280:30723–30734.

Schwarz DA, Katayama CD, Hedrick SM. 1998 Schlafen, a new family of growth regulatory genes that affect thymocyte development. Immunity 9:657–668.

Berger M, Krebs P, Crozat K, Li X, Croker BA, Siggs OM, Popkin D, Du X, Lawson BR, Theofilopoulos AN, et al. 2010 An Slfn2 mutation causes lymphoid and myeloid immunodeficiency due to loss of immune cell quiescence. Nat Immunol 11:335–343.

Patel VB, Yu Y, Das JK, Patel BB, Majumdar APN. 2009 Schlafen-3: a novel regulator of intestinal differentiation. Biochem Biophys Res Commun 388:752–756.

Li M, Kao E, Gao X, Sandig H, Limmer K, Pavon-Eternod M, Jones TE, Landry S, Pan T, Weitzman MD, et al. 2012 Codon-usage-based inhibition of HIV protein synthesis by human schlafen 11. Nature 491:125–128.

Ogura T, Wilkinson AJ. 2001 AAA+ superfamily ATPases: common structure--diverse function. Genes Cells 6:575–597.

Nedelsky NB, Todd PK, Taylor JP. 2008 Autophagy and the ubiquitin-proteasome system: collaborators in neuroprotection. Biochim Biophys Acta 1782:691–699.

Smith MH, Ploegh HL, Weissman JS. 2011 Road to ruin: targeting proteins for degradation in the endoplasmic reticulum. Science 334:1086–1090.

Varshavsky A. 2012 The ubiquitin system, an immense realm. Annu Rev Biochem 81:167–176.

Valastyan JS, Lindquist S. 2014 Mechanisms of protein-folding diseases at a glance. Dis Model Mech 7:9–14.

Hamilton AJ. 2010 MicroRNA in erythrocytes. Biochem Soc Trans 38:229–231.

Angénieux C, Maître B, Eckly A, Lanza F, Gachet C, de la Salle H. 2016 Time-Dependent Decay of mRNA and Ribosomal RNA during Platelet Aging and Its Correlation with Translation Activity. PLoS ONE 11:e0148064.

Qu BH, Strickland EH, Thomas PJ. 1997 Localization and suppression of a kinetic defect in cystic fibrosis transmembrane conductance regulator folding. J Biol Chem 272:15739–15744.

Futerman AH, van Meer G. 2004 The cell biology of lysosomal storage disorders. Nat Rev Mol Cell Biol 5:554–565.

Cox TM, Cachón-González MB. 2012 The cellular pathology of lysosomal diseases. J Pathol 226:241–254.

Pisarev AV, Unbehaun A, Hellen CUT, Pestova TV. 2007 Assembly and analysis of eukaryotic translation initiation complexes. Meth Enzymol 430:147–177.

